# Functional dissection of Leishmania major membrane components in resistance to cholesterol-dependent cytolysins

**DOI:** 10.64898/2025.12.12.693994

**Authors:** Chaitanya S. Haram, Sebastian Salinas, Coleman Wilson, Salma Waheed Sheikh, Kai Zhang, Peter A. Keyel

**Affiliations:** Department of Biological Sciences, Texas Tech University, Lubbock, TX 79409

**Keywords:** cholesterol-dependent cytolysins, streptolysin, perfringolysin, sterol accessibility, membrane defense, Leishmania major, glycans

## Abstract

Bacteria use cholesterol-dependent cytolysins (CDCs) to damage eukaryotes. While well-studied in mammals, the mechanisms by which CDCs bind to and kill protozoans remains unclear. CDCs bind to the human pathogen *Leishmania major*, but only kill in the absence of sphingolipids. The contribution of other leishmanial membrane components to CDC binding and cytotoxicity remains unknown. Here, we used genetic knockouts and inhibitors to determine the contribution of key membrane components to CDC binding and killing in *L. major*. We analyzed toxin binding and killing using flow cytometry and western blotting. Loss of the virulence factor GP63 enhanced toxicity of perfringolysin O, but not streptolysin O. Plasmenylethanolamine and lipophosphoglycan had minimal contributions to CDC binding and cytotoxicity. Removal of sterols protected cells from CDCs, yet failed to reduce binding. We used CDCs defective in engaging glycans or cholesterol to confirm that CDCs deficient in sterol binding, but not glycan binding, could bound to *L. major*. Thus, in non-mammalian systems, CDCs may rely on glycans for binding, while using sterols for pore-formation. This suggests that CDCs may not be sterol-specific probes in some non-mammalian systems. We conclude that early-branching eukaryotes use distinct mechanisms from mammals to limit CDC pore-formation and killing.

## INTRODUCTION

One neglected tropical disease with global impact is leishmaniasis. Leishmaniasis afflicts over two million, with 70,000 deaths annually, and is characterized by disfiguring skin or mucocutaneous lesions, and lethal visceral phenotypes [1]. Leishmaniasis is caused by deeply branching protozoan parasites in the genus *Leishmania*, including *L. major*. *L. major* cycles between mammalian hosts and sandfly vectors. In both life stages, *L. major* competes with bacteria and other microorganisms, though the competitive mechanisms remain poorly defined.

One aspect of interspecies competition is the secretion of and resistance to pore-forming toxins. Pore-forming toxins comprise the largest family of bacterial toxins [2], but their ability to destroy non-mammalian cells are poorly described. The best studied pore-forming toxins are cholesterol-dependent cytolysin (CDCs). CDCs are a family of β-barrel, pore-forming toxins that are secreted as monomers, and oligomerize to form 30 nm pores [3]. The archetypal CDCs streptolysin O (SLO) and perfringolysin O (PFO), from *Streptococcus pyogenes* and *Clostridium perfringens*, respectively, bind to cholesterol in the mammalian cell membrane via cholesterol recognition motifs (CRM) and then use cholesterol for pore-formation [4]. Differences in the cholesterol-binding region enables SLO to damage the membrane faster and with less sensitivity to the rest of the lipid environment compared to PFO, which takes longer to forms pores and has greater sensitivity to the surrounding environment [5, 6, 7]. The cholesterol binding domain of PFO is frequently used as a cholesterol sensor to determine the amounts of accessible cholesterol in mammalian membranes [8]. However, the binding of SLO and PFO to mammalian cell membranes might also involve glycans during initial monomer binding [9, 10]. Thus, CDCs are well-characterized in mammalian systems.

In contrast to mammalian systems, the mechanisms by which protozoans resist pore-forming toxins are poorly understood. While some protozoans prevent toxin binding, we previously showed that *L. major* is bound by CDCs [11]. However, loss of sphingolipids is necessary for killing [11]. We chose *L. major* as a system to understand how protozoans deal with pore-forming toxins because *L. major* must compete with bacteria that secrete pore-forming toxins in the sandfly [12], it is a pathogen with human clinical significance [1], and it is a genetically tractable system. Knockouts and complemented strains exist for many enzymes that synthesize key membrane components, including phospholipids, sphingolipids, ergosterol, and GPI-anchored molecules [13, 14, 15, 16, 17, 18]. Our prior work showed that inositol phosphorylceramide (IPC), the major sphingolipid in *L. major*, prevents CDC-mediated cytotoxicity, despite CDC binding [11]. The contributions of other membrane components, such as phospholipids, glycoconjugates, and GPI-anchored proteins, to CDC resistance remain unresolved.

In this study, we systematically examined the role of other *L. major* plasma membrane constituents—including GPI-anchored proteins, phospholipids, and sterols—in resistance to CDCs. Using genetically engineered *L. major* knockout strains and cytotoxicity assays with SLO and PFO, we assessed the contribution of specific membrane factors to toxin susceptibility. We found that the virulence factor lipophosphoglyan (LPG) and plasmalogen phospholipid plasmenylethanolamine (PME) are dispensable for protection. The GPI-anchored metalloproteinase GP63 conferred resistance to PFO, but not to SLO. Moreover, depletion of ergosterol protected cells from CDC-mediated killing, despite continued toxin binding. This binding is likely mediated by glycans because SLO variants deficient in sterol recognition still bound *Leishmania* membranes, while SLO deficient in glycan binding had worse recognition. These data suggest that in *Leishmania* spp., CDCs may use sterols for pore-formation instead of binding. Collectively, our findings indicate that *Leishmania* employ distinct, non-mammalian strategies to defend against CDCs, offering potential insights into novel therapeutic targets.

## RESULTS

### GPI anchored proteins may protect *L. major* from PFO

To determine the contribution of *L. major* plasma membrane components other than sphingolipids to CDC-binding and cytotoxicity, we first examined the abundant glycosylated virulence factors GP63 and LPG. Glycans may contribute to toxin binding in mammalian cells [9, 10], and our prior work with *spt2^—^* promastigotes suggested glycans contribute to toxin-binding in *L. major* [11]. To test the contribution of GP63 and LPG, we used promastigotes deficient in one of three genes: *gp63*, which encodes the GPI-anchored metalloprotease GP63 [18], *lpg1,* which encodes a galacto-furanosyltransferase pivotal for synthesis of the LPG glycan core [17] and *gpi8*, which encodes the transamidase that links GPI anchors to proteins [19].

Since the *gpi8^—^* did not exist in *L. major*, we created this mutant and complemented it via CRISPR (Supplementary Fig S1). To confirm the successful generation of the *gpi8^—^* mutant in *L. major*, promastigotes were screened for drug resistance and verified by PCR analysis. PCR amplification using *gpi8*-specific open reading frame (ORF) primers showed no detectable band in the *gpi8^—^*mutant, whereas amplification with selection marker–specific primers produced the expected fragments, confirming complete replacement of the *gpi8* alleles (Supplementary Figure S1A). In contrast, the wild type strain had the intact *gpi8* ORF and no signal for the drug maker. To verify genetic complementation, the *gpi8* ORF was episomally expressed in the *gpi8^—^*background. The presence of *gpi8* in the complemented line (*gpi8^—^/+*GPI8) was confirmed by PCR analysis (Supplementary Figure S1A). GPI8 is required for the synthesis of all GPI-anchored proteins. Consistent with loss of *gpi8*, the *gpi8^—^* mutant lost GP63 expression, whereas wild type and *gpi8^—^/+*GPI8 promastigotes maintained expression (Supplementary Figure S1B). Conversely, LPG was significantly elevated in the *gpi8^—^* mutant compared to wild type and *gpi8^—^/+*GPI8 promastigotes (Supplementary Figure S1B). These results validated the generation of *gpi8^—^* mutant line and genetic complementation of *gpi8^—^/+*GPI8, providing a reliable system to test the role of GPI-anchored proteins in *L. major*.

We then tested the binding and cytotoxicity of CDCs in *gp63^—^*, *lpg1^—^*, or *gpi8^—^* promastigotes, their wild type counterparts and complemented strains. The *gp63^—^* were generated in *L. major* strain Seidman, and are defective in GP63 synthesis [18], while the other strains were generated in strain LV39. Since our prior work in *spt2^—^* promastigotes [11] showed increased cytotoxicity despite no alterations to binding, we tested both binding and cytotoxicity of SLO and PFO. In the absence of the SPT2 inhibitor myriocin, SLO or PFO failed to kill any of the promastigotes (Supplementary Fig S2), consistent with the protection sphingolipids affords to *L. major* against CDCs [11]. When *gp63^—^*, *lpg1^—^*, or *gpi8^—^* promastigotes were treated with myriocin, CDC binding was similar across all groups (Fig 1A, B). These data indicate that CDCs can bind to *L. major* in the absence of glycans. Myriocin-treated *lpg1^—^* promastigotes showed equal levels of killing by SLO and PFO compared to myriocin-treated wild type (LV39) and *lpg1^—^*/+LPG1 cells (Fig 1C, F, Supplementary Fig S2A & B). Loss of GP63 trended to show a decrease in death by SLO (high LC_50_), which was not reversed in the complemented strain (Fig 1D, Supplementary Fig S2C). In contrast, loss of GP63 enhanced cytotoxicity against PFO, which was partially reversed in *gp63^—^*/+GP63 cells (Fig 1G, Supplementary Fig S2D). The myriocin-treated *gpi8^—^* and *gpi8^—^*/+GPI8 promastigotes largely phenocopied the *gp63^—^* and *gp63^—^*/+GP63 cells when challenged by SLO, including the failure of the *gpi8^—^*/+GPI8 to reverse the decreased killing (Fig 1E, Supplementary Fig S2E). Surprisingly, myriocin-treated *gpi8^—^* cells, which lack GP63 among other GPI-anchored proteins, were no more sensitive to PFO than wild type cells (Fig 1H, Supplementary Fig S2F). However, overexpression of GPI8 in these cells protected them from PFO, even at the supraphysiologic dose of 64,000 HU/mL (Fig 1H, Supplementary Fig S2F). Overall, we conclude that LPG fails to contribute to resistance to CDCs, while GP63 may protect *L. major* from PFO.

**Figure 1.**
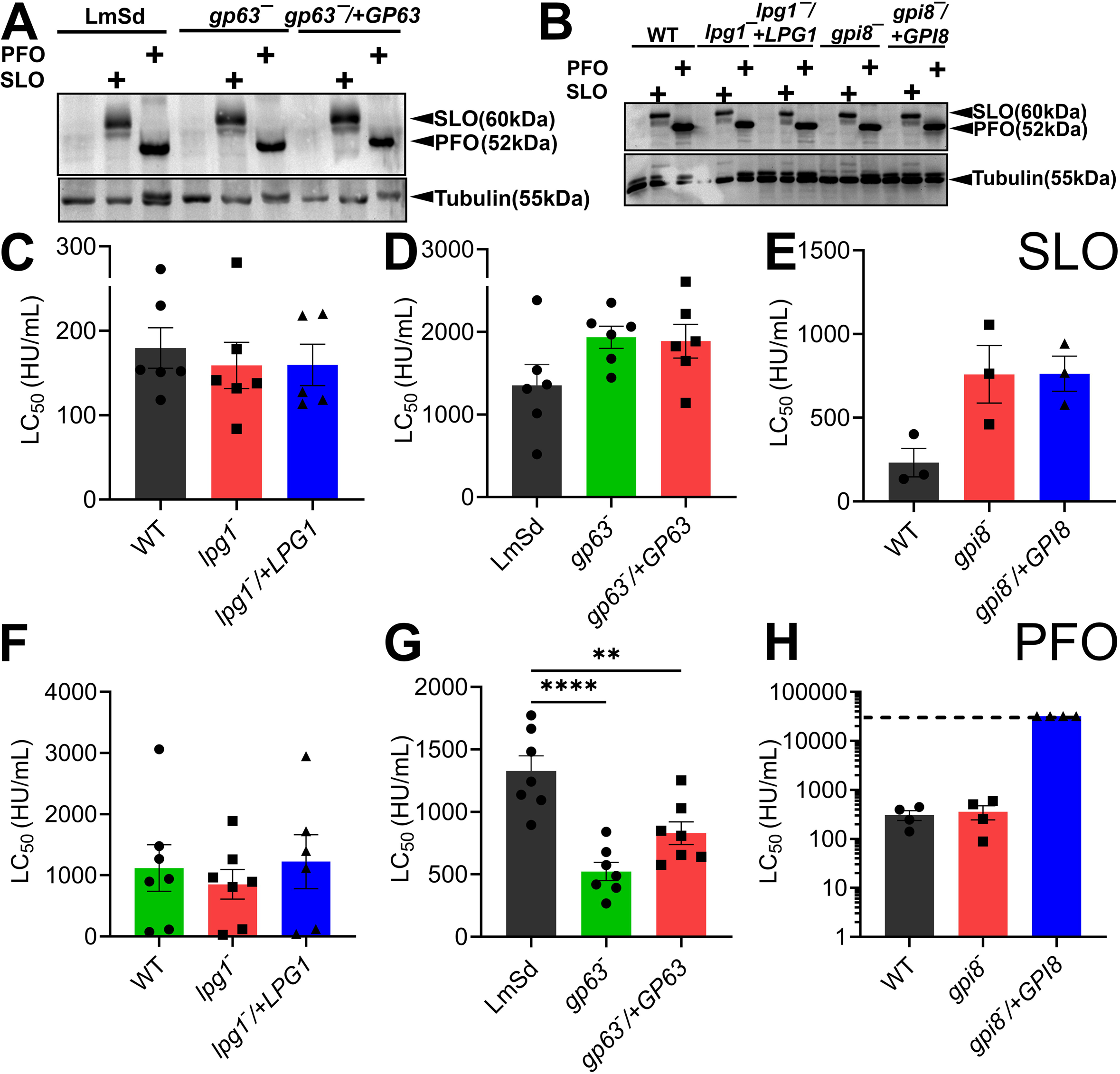
The metalloproteinase GP63 protects *L. major* from PFO. (A, B) LV39 wild type (WT), Seidman wild type (LmSd), *gp63^—^, gp63^—^*/+GP63, *lpg1^—^*, *lpg1^—^*/+LPG1, *gpi8^—^* & *gpi8^—^*/+GPI8 *L. major* promastigotes were challenged with 5 μg (A) SLO or (B) PFO at 4° C and then analyzed by western blotting using the indicated antibodies. (C-H) The indicated promastigotes were pretreated with 10 μM myriocin, and challenged with (C-E) SLO or (F-H) PFO at 37° C for 30 min. PI uptake was analyzed by flow cytometry. LC_50_ values were calculated via logistic modeling. The dashed line indicates the limit of detection (32,000 HU/mL). One representative blot out of 3 independent blots is shown. Graphs show the mean ± S.E.M. of (C, D, F, G) six, (E) three, or (F) four independent experiments, with each independent experiment plotted as a data point. ** p<0.01, **** p<0.0001 by one way ANOVA with multiple comparisons and Sidak-Bonferroni correction.

### Plasmenylethanolamine is unnecessary to resist CDCs

Kinetoplastids like *L. major* contain both standard phospholipids, plus plasmalogens like plasmenylethanolamine (PME). Plasmalogens contain an ether linkage at the sn1 and an ester linkage at the sn2 position of the phospholipid. PME comprises 80-90% of the ethanolamine-containing phospholipids in *L. major* promastigotes [16]. Genetic deletion of *ethanolamine phosphate cytidylyltransferase* (*ept*) eliminates plasmenylethanolamine from *L. major*, while phosphatidylethanolamine (PE) levels remain unperturbed due to production of PE from phosphatidylcholine (PC) [16]. PC is essential in *L. major* promastigotes and loss of its biosynthesis is lethal [20]. We tested the contribution of PME to toxin resistance by treating wild type, *ept*^—^, and *ept*^—^/+EPT promastigotes with myriocin and then measured binding and cytotoxicity. Myriocin pre-treated *ept*^—^ promastigotes bound SLO comparably to both wild type and *ept*^—^/+EPT promastigotes (Supplementary Fig. S3). We then challenged these cells with SLO or PFO. Without myriocin pre-treatment, all genotypes resisted both SLO and PFO (Supplementary Fig. S3). With myriocin pre-treatment, *ept*^—^ had equal sensitivity to both SLO and PFO compared to both wild type and *ept*^—^/+EPT promastigotes (Supplementary Fig. S3). Based on these data, we conclude that PME is dispensable for binding and cytotoxicity of SLO and PFO.

### Cholestane-based sterols reduce binding and cytotoxicity in *L. major*

We next tested the other major membrane component used by CDCs for binding and cytotoxicity: sterols. We used mutants lacking the key ergosterol synthesis genes *c14 demethylase* (*c14dm*) and *sterol 24-C methyltransferase* (*smt*). Instead of ergosterol, these mutants have elevated levels of 14-methylated ergosterol or cholestane-based sterols, respectively [15, 21]. The *c14dm^—^* promastigotes are more sensitive to membrane stress [15], including detergents like Triton-X-100 [11, 22] even without myriocin. We tested the binding of SLO to these cells. While SLO bound to *c14dm^—^* promastigotes like wild type, SLO bound less to the *smt^—^* promastigotes (Fig 2A). This reduced binding was rescued in the *smt^—^*/+SMT cells (Fig 2A). We then tested the cytotoxicity of CDCs in these cell types. Consistent with our prior work [11], these cells were resistant to CDCs without myriocin treatment (Supplementary Fig S4). In contrast to the increased detergent sensitivity, myriocin-treated *c14dm^—^* promastigotes showed equivalent sensitivity to both CDCs compared to wild type and complemented strains (Fig 2B, C, Supplementary Fig S4A, B). Consistent with the reduced binding, there was a trend towards resistance of the *smt^—^*cells to both CDCs that was reversed in the *smt^—^*/+SMT cells (Fig 2B, C, Supplementary Fig S4C, D). Based on these data, we conclude that CDCs do not bind well to cholestane-based sterols in the membrane, and this may reduce cytotoxicity of CDCs in *L. major*.

**Figure 2.**
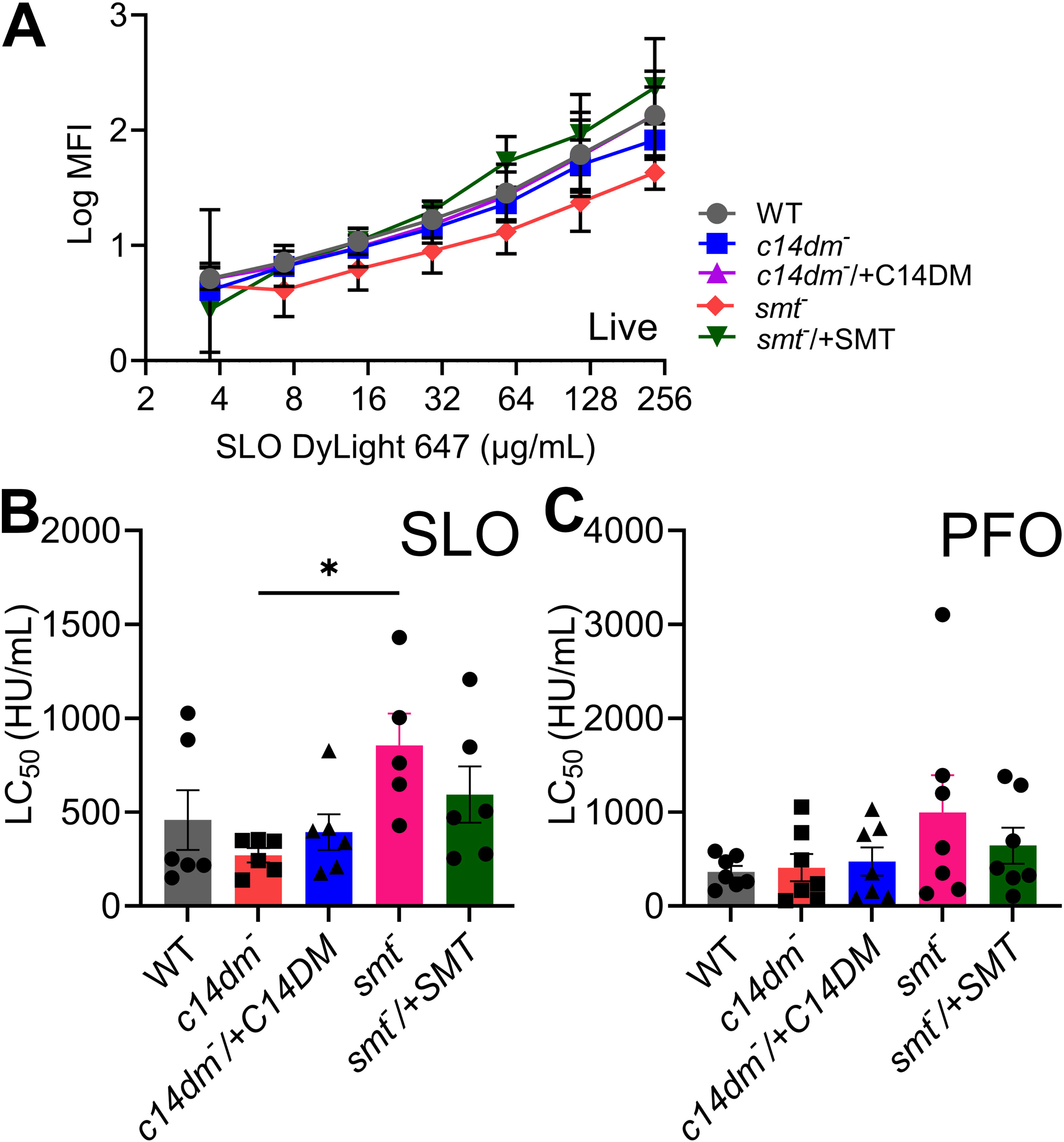
Cholestane based sterols limit binding and cytotoxicity of SLO but not PFO in *L. major* promastigotes. Wild type (WT), *c14dm^—^*, *c14dm^—^*/+C14DM, *smt^—^*, and *smt^—^*/+SMT *L. major* promastigotes pretreated with 10 μM myriocin were challenged with (A) SLO WT DyLight 647 (B) SLO, or (C) PFO. PI uptake and toxin binding was analyzed by flow cytometry. (A) Log median fluorescence intensity of DyLight 647 fluorescence gated on live cells is shown. LC_50_ values were calculated via logistic modeling. Graphs display mean ± SEM of (A) three, (B) six, or (C) seven independent experiments, with (B, C) independent experiments plotted as data points. *p<0.05, **p<0.01 by 2 way ANOVA with multiple comparison and Sidak-Bonferroni correction.

### Ergosterol is needed for CDC cytotoxicity but not binding in promastigotes

We further explored sterol dependence using chemical approaches to probe the contribution of sterols. We previously showed that *spt2^—^*cells are more sensitive than wild type cells to Triton-X-100 [11]. We extended this work by testing their sensitivity to the sterol-binding detergent saponin. The *spt2*^—^ promastigotes were more sensitive to saponin than wild type or *spt2*^—^ /+SPT (Supplementary Fig S5A).

We next used a less invasive method to remove membrane sterols: short-term treatment with 2-hydroxypropyl-β-cyclodextrin (HPCD). In mammalian cells, short-term treatment removes accessible cholesterol from the plasma membrane without depleting intracellular stores [8, 23]. We treated HeLa cells as a control, or wild type, *spt2*^—^, or *spt2*^—^ /+SPT promastigotes with HPCD, then challenged them with SLO or PFO, and measured binding and cytotoxicity. HPCD pretreatment decreased toxin binding to HeLa cells, but failed to reduce binding to *L. major* promastigotes (Fig 3). Indeed, quantitation of the blots revealed a slight trend towards increased binding of SLO and PFO after HPCD treatment in *L. major* promastigotes (Fig 3). Overall, limiting ergosterol in the plasma membrane using HPCD failed to reduce CDC binding in *L. major*.

**Figure 3.**
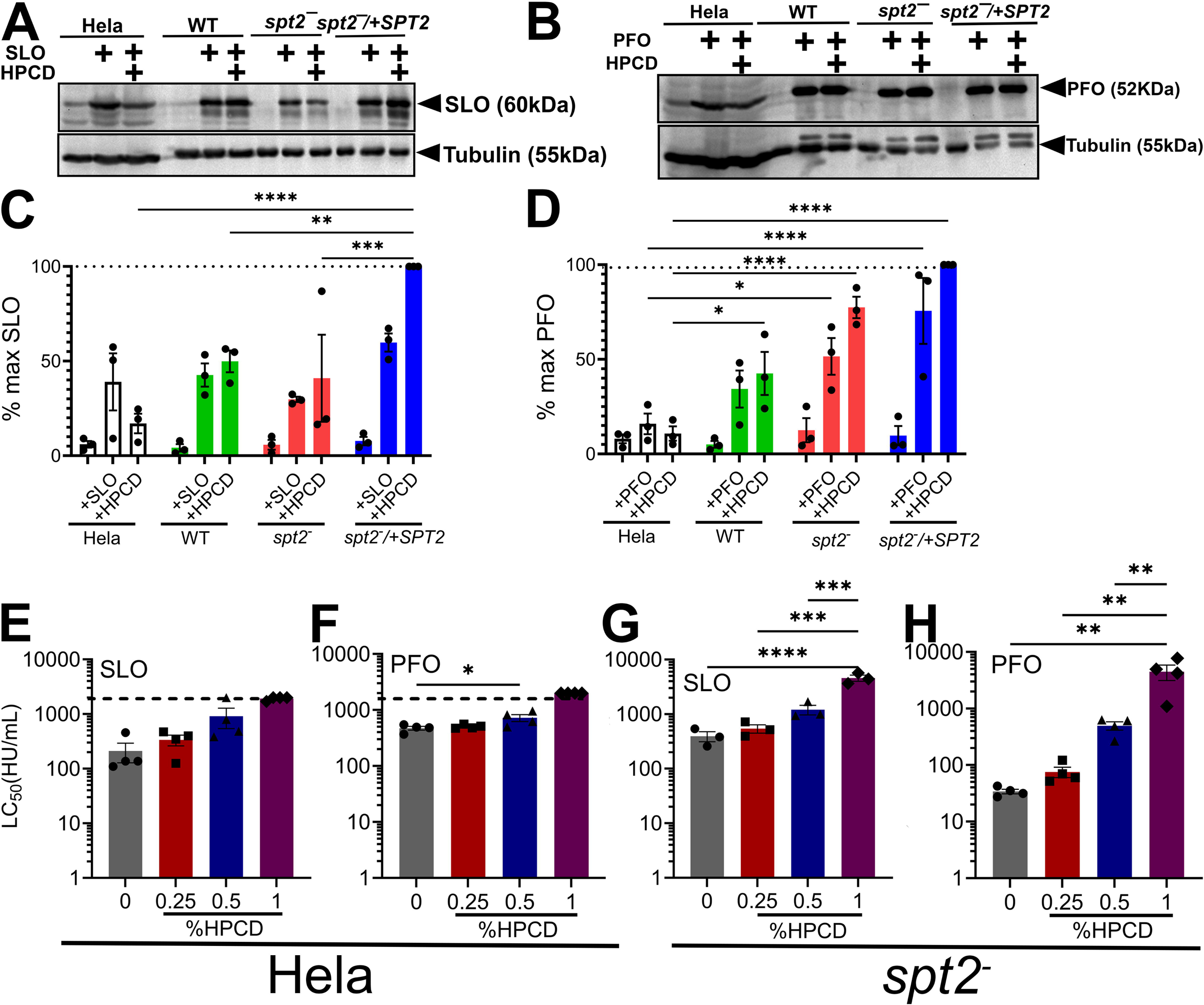
CDC cytotoxicity is ergosterol-dependent in *L. major*. (A-D) HeLa cells, wild type (WT), *spt2^—^*, or *spt2^—^*/+SPT2 *L. major* promastigotes pretreated with 1% HPCD were challenged with 5 μg (A, C) monomer locked SLO or (B, D) monomer locked PFO at 4° C for 30 min. Cells were analyzed by western blotting for SLO, PFO and tubulin. (C, D) Protein levels were quantitated by densitometry. (E-H) spt2*^—^ L. major* promastigotes or Hela cells pretreated with the indicated concentrations of HPCD were challenged with SLO or PFO for 30 min at 37°C. PI uptake was analyzed by flow cytometry. LC_50_ values were calculated via logistic modeling. Graphs display mean ± SEM of (G) three or (E,F, H) four independent experiments, with (E-H) independent experiments plotted as individual points. Representative blots from three independent experiments are shown. Blots were normalized to the greatest CDC expression, which is indicated by the dashed line. *p<0.05, **p<0.01, ***p<0.001 by one way ANOVA with multiple comparison and Sidak-Bonferroni correction.

We next tested cytotoxicity in HPCD-treated cells. We observed a HPCD dose dependent increase in resistance to SLO and PFO of *spt2*^—^ cells (Fig 3, Supplementary Fig S5B, C), which was consistent with an HPCD dose-dependent increase in resistance to SLO and PFO in Hela cells (Fig 3, Supplementary Fig S5D, E). We conclude that sterols are needed for cytotoxicity, but not for the binding of CDCs to *L. major*.

### SLO uses glycan binding residues to bind *L. major* promastigotes

Since CDCs bound to *L. major* independent of sterol removal, we tested the contribution of the SLO sterol and glycan binding sites to *L. major* binding. SLO sterol binding in mammalian cells can be abrogated by the SLO ΔCRM mutant [4], while glycan binding is lost in the SLO Q476N mutant [10]. We previously showed that neither of these mutant toxins kill *L. major* [11], but we did not test binding. We measured the binding of wild type SLO, SLO Q476N or SLO ΔCRM to HeLa cells, wild type, *spt2*^—^, or *spt2*^—^ /+SPT promastigotes with and without HPCD. Consistent with previous results [10], both SLO Q476N and ΔCRM had minimal binding to Hela cells independent of HPCD (Fig 4). In *L. major*, both SLO Q476N and ΔCRM had reduced binding independent of genotype, with SLO ΔCRM trending towards greater binding (Fig 4). The weak binding from SLO Q476N showed a trend to further reduction after HPCD treatment in WT and *spt2*^—^ /+SPT (Fig 4)., Together, we interpret these results to indicate that the glycan binding site is important for binding to *L. major*, with the sterol engagement more important for toxicity.

**Figure 4.**
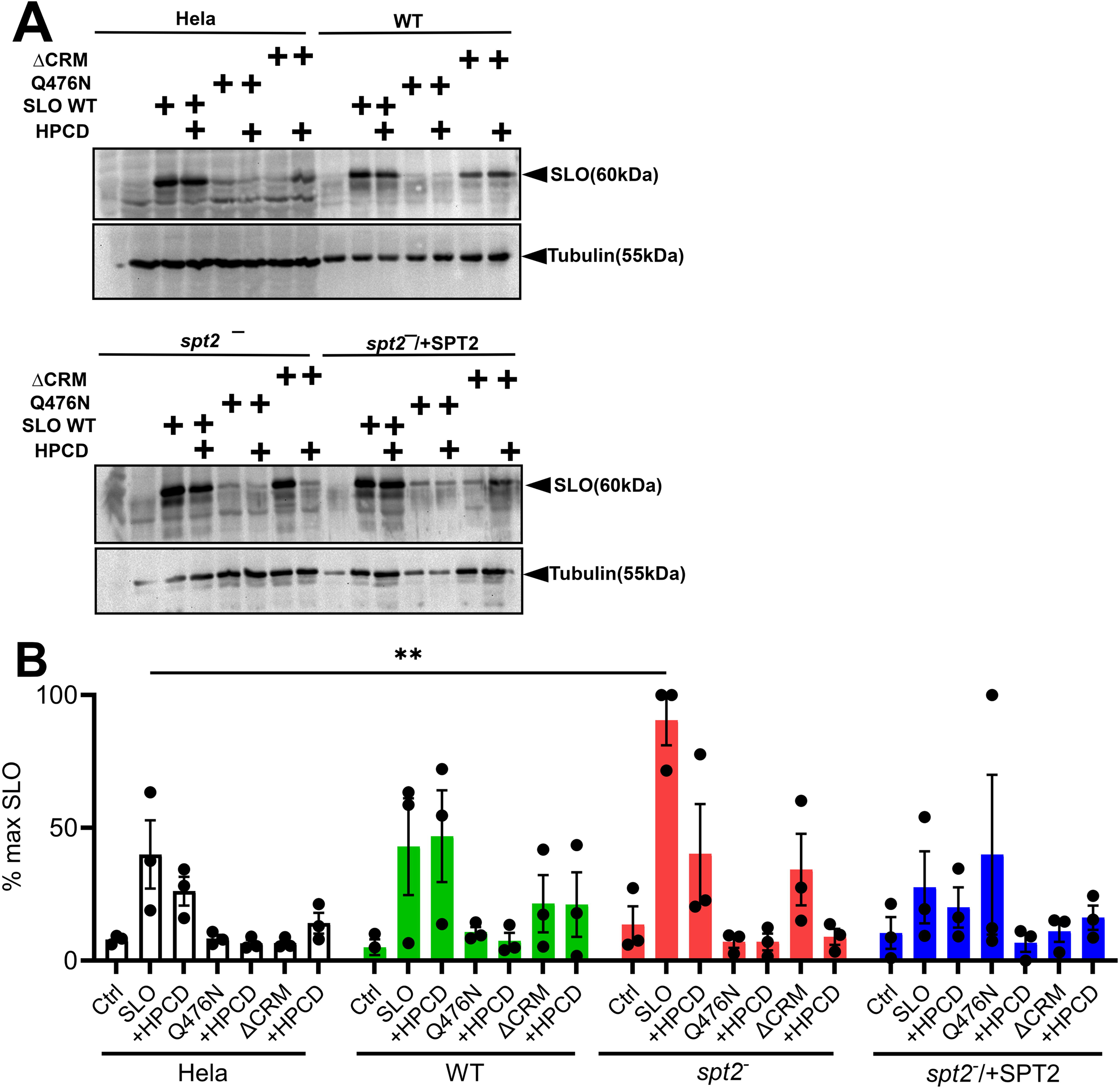
SLO glycan binding determinants are pivotal for binding to *Leishmania major* promastigotes. Hela cells, wild type (LV39WT), *spt2*^—^, or *spt2*^—^/+SPT2 *L. major* promastigotes were pretreated with 1% 2-hydroxypropylcyclodextrin (HPCD) at 27° C for 30 min, challenged with 5 μg wild type SLO (SLO WT), SLO Q476N or SLO ΔCRM at 4° C for 30 min and lysed for western blot analysis. Blots were probed with antibodies against SLO or tubulin. Representative blots from three independent experiments are shown. Blots were normalized to the greatest SLO expression in each experiment. **p<0.01 by one way ANOVA with multiple comparison and Sidak-Bonferroni correction.

## Discussion

In this study, we tested the contribution of leishmanial plasma membrane components to protection from CDCs. We found that GPI-anchored proteins may contribute to protection against PFO, while PME and LPG failed to limit CDC cytotoxicity. Cholestane-based sterols reduced CDC binding. Removal of ergosterol from the plasma membrane limited CDC cytotoxicity without reducing binding. Consistent with this finding, SLO lacking the cholesterol recognition motif could bind, but SLO deficient in glycan binding failed to bind. We interpret these data to suggest that single-celled protozoans like *Leishmania* use distinct defense strategies compared to mammalian cells.

Kinetoplastids rely on the packing of GPI-anchored proteins and glycans to protect them from environmental and immune stressors. We found that in *L. major*, loss of GP63 sensitized cells to PFO while a global loss of GPI-anchored proteins failed to alter cellular sensitivity. However, overexpression of GPI8 made the cells resistant to PFO. Since binding was not altered, it is possible that GP63 and other GPI-anchored proteins interfere with oligomerization or pore formation. GP63 localizes to the detergent-resistant membranes (DRMs) enriched in sterols, so the tight packing of GP63 could limit PFO cytotoxicity [24]. PFO has stricter membrane binding requirements than SLO in both mammals and *L. major* [5, 11], which may account for why SLO is not affected by GP63, but PFO is. LPG and PME localization to the non-DRM fraction [16, 25] could account for the failure of these species to protect cells from CDC cytotoxicity. Glycan-preference could also account for differences in cytotoxicity. In mammalian systems, SLO prefers galactose-rich glycans, while PFO prefers N-acetylneuraminic acid in glycans [9, 10]. In *L. major*, LPG is rich in galactose residues interspersed with mannose residues, while GP63 is mannose-rich [26]. Thus, these results provide new insight into *L. major* membrane environment CDCs, especially PFO, require for cytotoxicity.

The key lipid for CDC activity is sterol. In mammalian cells, SLO and PFO use cholesterol both for binding and pore-formation [2, 4], while other CDCs rely on the GPI-anchored protein CD59 for binding and cholesterol for pore formation [27]. We found that the accumulation of cholestane-based sterols in the *smt^—^* mutant [14] reduced both binding and cytotoxicity of SLO in *L. major*. In contrast, methylation on C24 made no difference for CDC cytotoxicity. This is surprising because *c14dm^—^* promastigotes are generally more sensitive to membrane stress [15]. Whether these differences are due to lipid packing, or direct engagement of the CDC with the sterol remain to be determined.

In contrast to other systems, where cholesterol can be sufficient for CDC binding and cytotoxicity [2], we found that ergosterol is only needed for cytotoxicity, not binding in *L. major* promastigotes. These findings are consistent with previous reports in mammalian cells that SLO and PFO can bind to glycans and or glycosphingolipids in addition to cholesterol, while the process of membrane insertion depends on cholesterol [9, 10]. We used toxins defective in cholesterol or glycan binding in the presence of HPCD to tease apart the relative contributions of both sites. While both mutants had reduced binding compared to wild type, they bound better than to HeLa cells. This suggests the CDCs may use the CRM to recognize membrane components other than ergosterol in the leishmanial membrane. This is consistent with the failure of HPCD to eliminate binding of the glycan-binding mutant. Overall, we find that ergosterol is needed for CDC-mediated killing, but the CDCs use both glycan-binding and cholesterol-recognition regions to bind to multiple cryptic sites on the promastigote membrane.

Our findings provide new insights into the competition between *L. major* and bacteria. Promastigotes compete with bacteria in the sandfly midgut. While IPC is the major protective determinant against CDCs in *L. major*, the toxins still bind. The recent discovery of CDC-like toxins in *Elizabethkingia anopheles* and other bacteria [28], suggests that bacterial pore-forming toxins are used for more than human pathogenesis. While it is possible that toxin binding to *L. major* is incidental to their major function against other organisms, our work raises the possibility that these toxins could have secondary roles opsonizing *L. major*. Notably, SLO is capable of translocating the lethal NAD+ glycohydrolase (Spn) across mammalian membranes via its N-terminal domain [29]. We did not test Spn in *L. major*, but that could be a back-up killing mechanism bacteria use to compete. *Clostridium perfringens* α-toxin has sphingomyelinase activity [30], which suggests PFO could be lethal if α-toxin cleaves IPC. Future work is needed to determine how CDC binding alters the balance between *L. major* and the sandfly gut microbiota.

While this study provides a comprehensive analysis of key *Leishmania major* plasma membrane components involved in defense against pore-forming toxins, it had some limitations. We did not test amastigotes because *L. major* amastigotes cannot be grown axenically. Purification from macrophages would make killing and binding assays challenging to interpret. We only investigated one phospholipid, PME. The functional roles of other abundant phospholipids, such as PC and PE remain unexplored because their genetic knockout is lethal. Differences in our findings between *gpi8*^—^ and *gp63*^—^ promastigotes suggest that the minor GPI-anchored molecules contribute to binding and cytotoxicity. However, identifying which of components contribute is beyond the scope of our study. Due to this, we were unable to pinpoint the specific glycans in *L. major* promastigotes responsible for CDC binding. The present work provides a foundation for future studies to address these gaps and better understand membrane integrity and repair in *L. major*.

## Materials and Methods

### Reagents

All reagents were from Thermofisher Scientific (Waltham, MA, USA), unless otherwise noted. Cysteine-less His-tagged PFO (PFO WT) in pET22 was a generous gift from Rodney Tweten (University of Oklahoma Health Sciences Center, Oklahoma City, OK, USA). Cysteine-less, codon-optimized SLO (SLO WT) in pBAD-gIII, monomer-locked (G298V/G299V) PFO, monomer-locked (G398V/G399V) SLO, SLO S101C, SLO Q476N and SLO ΔCRM (T564A/L565A were previously described [11]. 6D11 anti-SLO monoclonal antibody (mAb) was from Fisher (catalog NBP105126). WIC79.3 anti-LPG mAb was from Stephen Beverley (Washington University School of Medicine, St Louis, MO, USA) [31]. Anti-GP63 monoclonal antibody m235 was from Robert McMaster (The University of British Columbia, Vancouver, BC, Canada) [32]. Mouse anti-tubulin antisera was from Fisher. Cross-adsorbed HRP-conjugated anti-mouse IgG or anti-rabbit IgG was from Jackson Immunoresearch (West Grove, PA).

### Recombinant Toxins

Toxins were induced and purified as previously described [33, 34]. Toxins were induced with 0.2% arabinose (SLO), or 0.2 mM IPTG (PFO) for 3 h at room temperature and purified using Nickel-NTA beads. Protein concentration was determined by Bradford assay and hemolytic activity was determined as previously described using human red blood cells (Zen Bio, Research Triangle Park, NC, USA) [34]. One hemolytic unit is defined as the amount of toxin required to lyse 50% of a 2% human red blood cell solution in 30 min at 37 °C in 2 mM CaCl_2_, 10 mM HEPES, pH 7.4, and 0.3% BSA in PBS. Hemolytic units were used to control for differences in toxin activity. The specific activities of monomer-locked toxins, SLO Q476N and SLO ΔCRM were <100 HU/mg. They were used at a mass equivalent to wild type SLO and PFO. Multiple toxin preparations were used to control for variation by preparation (Table 1). Where indicated, wild type or monomer-locked toxins were conjugated to Cy5 using NHS chemistry as described [33]. Alternatively, SLO S101C, which retains wild type activity [33], was conjugated to AlexaFluor647 or DyLight 647 using maleimide chemistry as described [33].

**Table 1.**
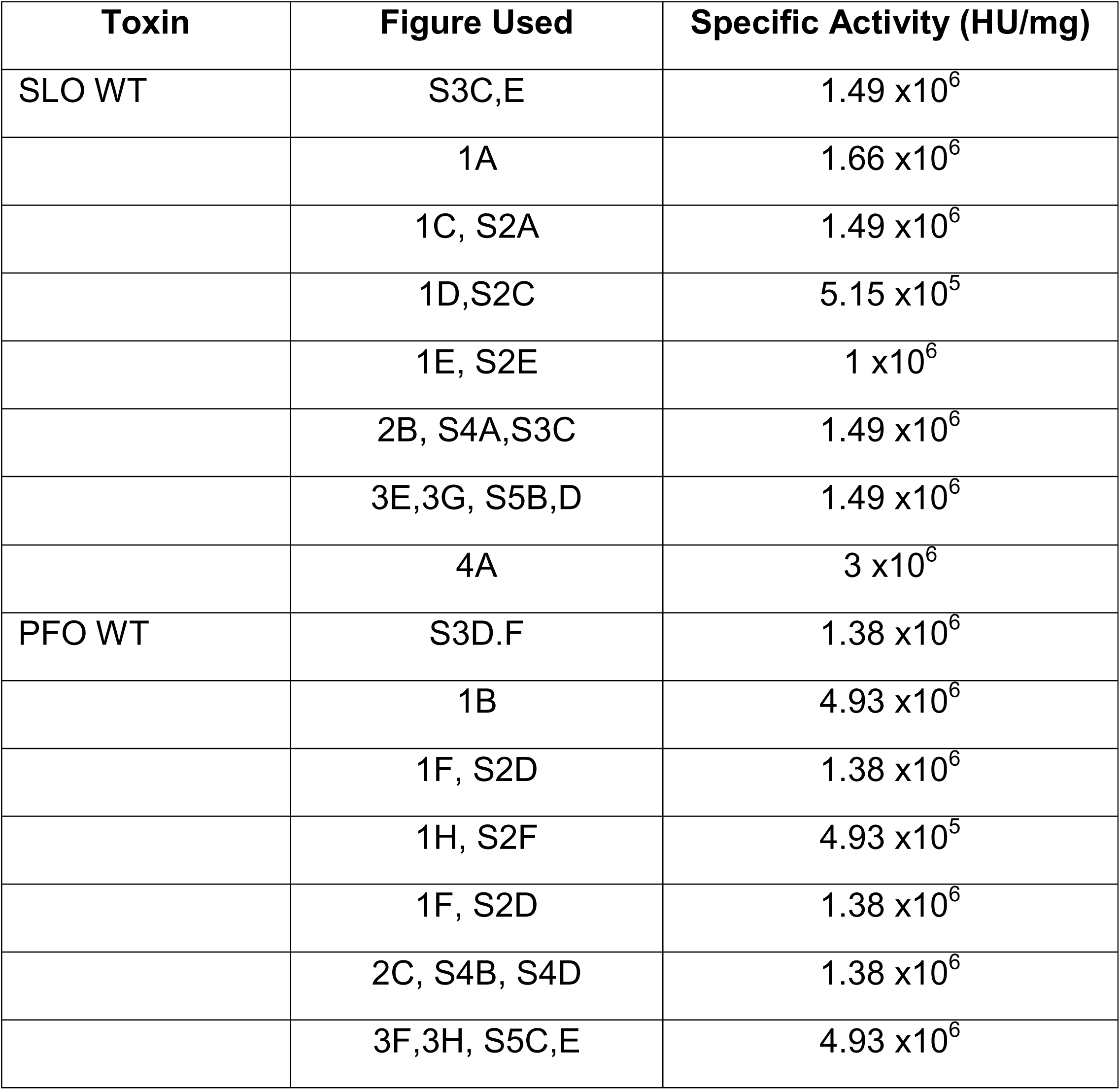
Specific activity of active toxin preps used.

### Leishmania strains and culture

Leishmania major strains and mutants were previously characterized (Table 2). LV39 clone 5 (Rho/SU/59/P) was used as the wild type strain for all experiments except when the *gp63*^—^ *L. major* was used. For these experiments, the *L. major* strain NIH S (MHOM/SN/74/Seidman) clone A2 was used as the wild type strain [18]. *L. major* promastigotes were cultured at 27°C in 1XM199 medium (pH 7.4) with 10% fetal bovine serum and additional supplements. The *gpi8^—^* mutant was generated using the CRISPR-Cas9 system as described [35]. The endogenous *gpi8* gene was disrupted by replacing *gpi8* with a drug-resistance marker through homologous recombination. The resulting *gpi8^—^* was selected based on resistance to blasticidin (BSD) and hygromycin (HYG). The replacement of *gpi8* alleles was confirmed by PCR using ORF primers and drug specific primers to confirm integration. For genetic complementation, *gpi8* was cloned into pXG1a. The construct was then transfected into *gpi8^—^* promastigotes to generate the *gpi8^—^*/*+*GPI8 complemented strain.

**Table 2.** Leishmania strains used in this study.

| Name | Genotype | Background | Reference |
| --- | --- | --- | --- |
| *spt2*<sup>−</sup> | Δ*spt2*::HYG/Δ*spt2*::PAC | LV39 | [13] |
| *spt2*<sup>−</sup>/+SPT2 | Δ*spt2*::HYG/Δ*spt2*::PAC/+pXG-SPT2 | LV39 | [13] |
| *smt*<sup>−</sup> | Δ*smt*::HYG/Δ*smt*::PAC | LV39 | [21] |
| *smt*<sup>−</sup>/+SMT | Δ*smt*::HYG/Δ*smt*::PAC/+pXG-SMT | LV39 | [21] |
| *c14dm*<sup>−</sup> | Δ*c14dm*::HYG/Δ*c14dm*::PAC | LV39 | [15] |
| *c14dm*<sup>−</sup>/+C14DM | Δ*c14dm*::HYG/Δ*c14dm*::PAC/+pXG-C14DM | LV39 | [15] |
| *ept*<sup>−</sup> | Δ*EPT*::SAT/Δ*EPT*::HYG | LV39 | [16] |
| *ept*<sup>−</sup>/+EPT | Δ*EPT*::SAT/Δ*EPT*::HYG/1pXG1- EPT | LV39 | [16] |
| *lpg*<sup>−</sup> | Δ*LPG1*::HYG/Δ*LPG1*::PAC | LV39 | [17] |
| *lpg*<sup>−</sup>/+LPG | Δ*LPG1*::HYG/Δ*LPG1*::PAC/+pSNBR-LPG1 | LV39 | [17] |
| *gp63*<sup>−</sup> | Δ*GP63*::SAT/Δ*GP63*::HYR | Seidman | [18] |
| *gp63*<sup>−</sup>/+GP63 | Δ*GP63*::SAT/Δ*GP63*::HYR/+ pLEXNeo-gp63 | Seidman | [18] |
| *gpi8*<sup>−</sup> | Δ*GPI8*:: BSD/ Δ*GPI8*:::HYR | LV39 | This study |
| *gpi8*<sup>−</sup>/+GPI8 | Δ*GPI8*:: BSD/ Δ*GPI8*:::HYR/+ pXG1a-gpi8 | LV39 | This study |

Promastigotes were cultured at 27°C in M199 medium (Gibco, Waltham, MA) with 0.182% NaHCO_3_, 40 mM HEPES, pH 7.4, 0.1 M adenine, 1 µg/mL biotin, 5 µg/mL hemin, 2 µg/mL biopterin and 10% heat inactivated fetal bovine serum, pH 7.4. Episomal complemented cells were maintained in the same medium with the addition of 10 µg/mL neomycin (G418) except for experimental passages. Culture density and cell viability were determined by hemocytometer counting and flow cytometry after propidium iodide (PI) staining at a final concentration of 20 µg/mL. Log phase promastigotes were replicative parasites at 2.0 – 8.0 ×10^6^ cells/mL.

### HeLa cell culture

Hela cells (ATCC (Manassas, VA, USA) CCL-2) were maintained at 37°C, 5% CO_2_ in DMEM (Corning, Corning, NY, USA) supplemented with 10% fetal bovine serum (R&D Biosystems, Minneapolis, MN, USA) and 1× L-glutamine (D10). They were negative for mycoplasma by microscopy.

### Myriocin treatment of L. major

Log phase cells were seeded at 1.0 ×10^5^ cells/mL in complete medium and either treated with 10 µM Myriocin dissolved in 1X DMSO (experimental) or an equivalent volume of diluent 1x DMSO (control). Cells were cultured and allowed to reach log phase in 48 hours before harvesting and processing cells for experiments.

### Leishmania processing

Cells were processed as described [11] and either resuspended in resuspended in serum free 1X M199 to a final concentration of 1.0 X10^7^ cells/mL for Western blot or in serum free Tyrode’s buffer (137 mM NaCl, 2.7 mM KCl, 1.8 mM CaCl_2_, 0.49 mM MgCl_2_, 0.42 mM NaH_2_PO_4_, 11.9 mM NaHCO_3_, and 11.1 mM dextrose, pH 7.5) to a final concentration of 1.0 ×10^6^ cells/mL for cytotoxicity. For cytotoxicity, 1.0 ×10^5^ cells were plated per well in a V-bottom 96 well plate or per Marsh tube.

### Binding assay via western blot

*L. major* promastigotes were resuspended to a final cell concentration of 1.0X10^7^ cells in serum free 1X M199 supplemented with 2 mM CaCl_2_. Where indicated, *L. major* promastigotes were pretreated with 0%, 0.25%, 0.5%, or 1% HPCD for 30 min at 27° C before toxin challenge. Cells were challenged with 5 μg of SLO WT, PFO WT, SLO ML, PFO ML, SLO ΔCRM, or SLO Q476N for 30 min at 4° C. After toxin challenge, cells were centrifuged for 10 min at 3000 RPM (Rotor SX 4750) at room temperature (25° C). Cells were washed once with 1X PBS, centrifuged at 3000 RPM for 10 min at room temperature (25° C). Cell pellets were washboarded and then resuspended in 1X SDS sample buffer (with freshly added 2-mercaptoethanol) and heat at 95° C for 10 min. SDS-PAGE and western blots were performed as previously described [36]. Cell lysates were resolved on 10% acrylamide gels, and transferred to nitrocellulose at 110 V for 90 min. Blots were blocked with 5% bovine serum albumin (BSA) in 1X Tris-buffered saline (15.2mM NaCl, Tris base 46.2mM, 150 mM NaCl) with 0.1% Tween (TBST) at 4° C for 2 h, probed with primary antibodies overnight in 1% BSA in 1x TBST at 4^°^C, washed thrice with 1X TBST for 10 min each, incubated with HRP-conjugated secondary antibodies in 1% BSA in 1X TBST for 1 h, washed with 1X TBST thrice for 10 min each, developed with enhanced chemiluminescence reagent (1.25 mM luminol (Sigma), 0.01% H_2_O_2_ (Walmart, Fayetteville, AR), 0.2 mM p-coumaric acid (Sigma), 0.1 mM Tris-HCl, pH 8.4) and imaged on an iBlot (Invitrogen). Antibody dilutions were anti-SLO 1:2000, anti-LPG 1:1000, anti-GP63 1:1000, tubulin 1:1000, anti-mouse HRP 1:10,000, and anti-rabbit HRP 1:10,000.

### Protein expression levels of GPI anchored proteins

Total protein lysates were prepared from promastigotes of LV39WT, LV39Cas9, *gpi8^—^*, or *gpi8^—^*/*+*GPI8 promastigotes. Parasites were harvested at late-log phase, washed in PBS, and lysed in SDS containing buffer by boiling at 95°C for 10 min. Equal amounts of protein were resolved using SDS-PAGE gels and transferred to PVDF membranes. Membranes were blocked with 5% BSA in TBST and incubated with anti-GP63 (1:1000), anti-LPG WIC79.3 (1:1000), or anti-tubulin (1:1000) primary antibodies. HRP-conjugated anti-mouse antibodies were used for detection, and signals were visualized by chemiluminescence. Parallel blots were stained with Ponceau S staining solution to verify equal protein loading across samples.

### Flow cytometry cytotoxicity assay

Killing assays were performed as described [36]. Promastigotes were resuspended at 1 ×10^6^ cells/mL in Tyrode’s buffer supplemented with 2 mM CaCl_2_ and 20 μg/mL PI. HeLa cells were resuspended at 1 ×10^6^ cells/mL in RPMI media supplemented with 2 mM CaCl_2_ and 20 μg/mL PI. When indicated, cells were treated with 0.25%, 0.5% or 1% HPCD at 27° C for 30 min. Cells were challenged with toxins for 30 min at 37 °C and analyzed using an Attune flow cytometer. Specific lysis and LC_50_ were determined as described [36]. For fluorescence binding assays, Cy5, Alexa Fluor 647, or DyLight 647 conjugated toxins were used. The median fluorescence intensity (MFI) was quantitated, and background-subtracted using cells receiving no fluorescent toxin.

### Statistics

Prism (Graphpad, San Diego, CA), Sigmaplot 11.0 (Systat Software Inc, San Jose, CA) or Excel were used for statistical analysis. Data are represented as mean ± SEM. The LC_50_ for toxins was calculated by logistic modeling [36]. Statistical significance was determined by one-way ANOVA with Tukey post-testing, one-way ANOVA (Brown-Forsythe method) with Dunnett T3 post-testing, or Kruskal-Wallis, as appropriate. p < 0.05 was considered statistically significant. Graphs were generated in GraphPad and organized in Photoshop (Adobe, San Jose, CA, USA).

## Author Contributions

Conceptualization, CSH and PAK.; methodology, CSH, SS, CW, SWS.; formal analysis, CSH, SS, CW, PAK.; investigation, CSH, SS, CW, SWS.; resources, KZ; data curation, CSH, SS, CW, SWS, PAK.; writing^—^original draft preparation, CSH, SS, CW, SWS, PAK.; writing^—^review and editing, CSH, SS, CW, SWS, KZ, PAK..; visualization, CSH, SWS.; supervision, KZ, PAK.; project administration, KZ, PAK; funding acquisition, PAK. All authors have read and agreed to the published version of the manuscript.

## Funding

This work was supported by the National Institute of Allergy and Infectious Diseases of the National Institutes of Health grant R21AI156225 (PAK).

## Institutional Review Board Statement

Not applicable

## Data Availability Statement

All data are in the main text or supplementary materials.

## Acknowledgments

The authors thank the Keyel lab members for their critical review of the manuscript and the College of Arts & Sciences Microscopy for using their facilities.

## Conflicts of Interest

PAK is a co-founder of Ardiyon Bio. The funders had no role in the study’s design, data collection, analysis, or interpretation, manuscript writing, or decision to publish the results.

