## Supplemental Figures S1-S5 for "Functional dissection of Leishmania major membrane components in resistance to cholesterol-dependent cytolysins"

Supplementary Figures S1-S5

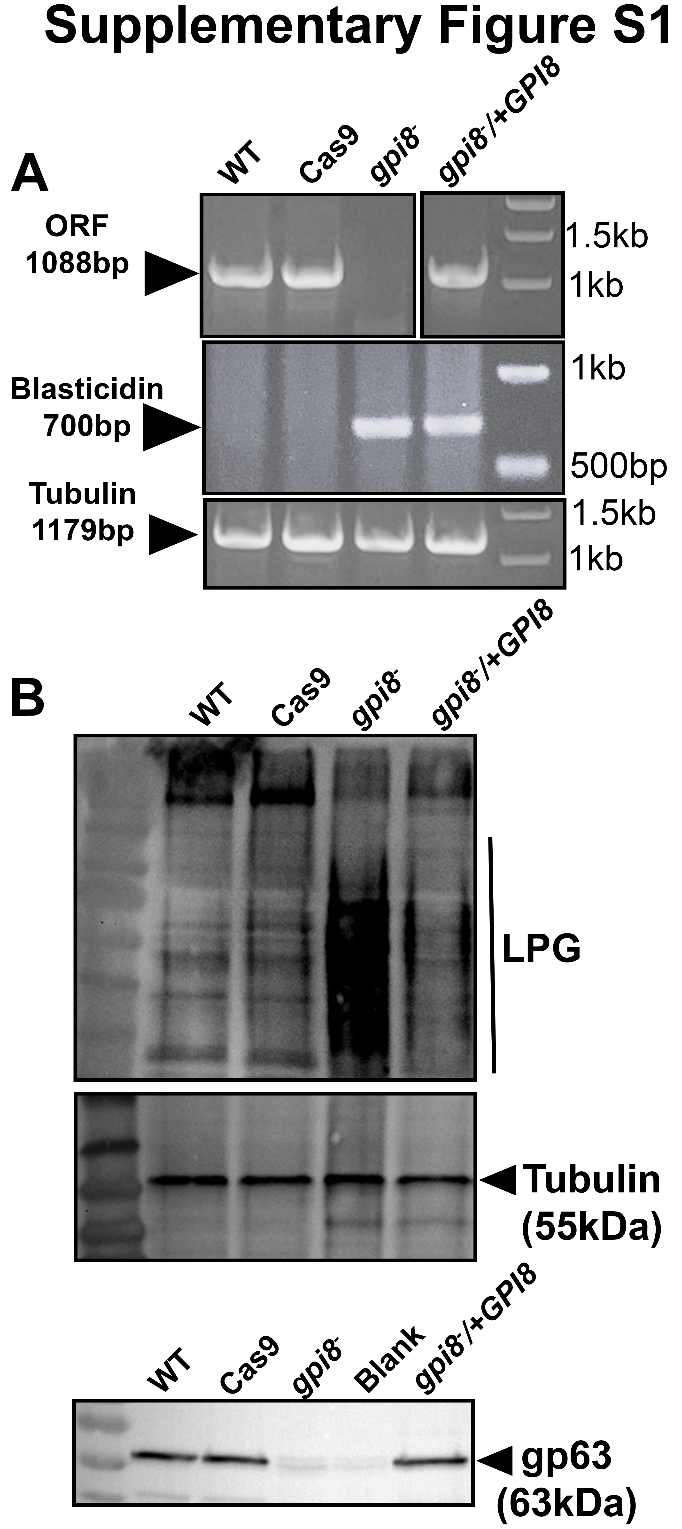

**Supplementary Figure S1.** **Generation of the *gpi8*^—^ and *gpi8*^—^/+GPI8 *L. major* strains.** (A) Genomic DNA was extracted after 10-15 days of transfection from LV39WT, LV39 Cas9, *gpi8^—^* and *gpi8^—^*/+GPI8 promastigotes and analyzed by PCR using primers for the open reading frame (1088bp), Blastidicin (700bp) or alpha tubulin (1179bp) on 1% agarose gel. (B-D) Total protein lysates were used to validate the functionality of *gpi8*. Blots were probed with the indicated antibodies. Blot is representative of 3 independent blots.

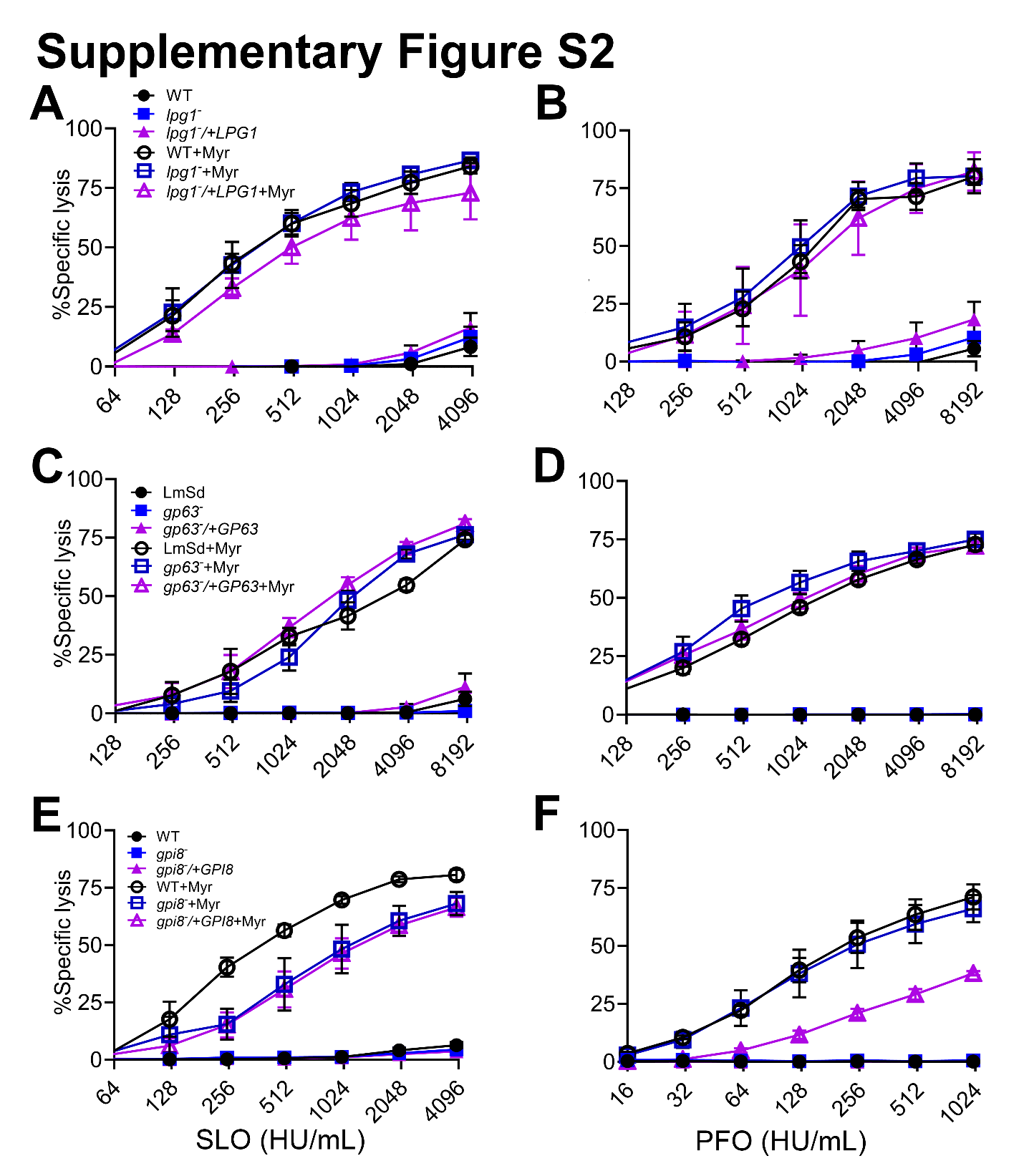

**Supplementary Figure** **S2. GPI null mutants of *L. major* are resistant to CDC cytotoxicity.** LV39 wild type (LV39WT), Seidman wild type (WT (LmSd), *lpg1^—^*, *lpg1^—^*/+LPG1, *gp63^—^, gp63^—^*/+GP63, *gpi8^—^* or *gpi8^—^*/+GPI8 *L. major* promastigotes grown in 1X M199 supplemented with either no myriocin, or 10 μM myriocin (+Myr) were challenged with (A,C,E) SLO or (B,D,F) PFO at 37° C for 30 min. PI uptake was measured by flow cytometry. Graphs display mean ± SEM of 3 independent experiments.

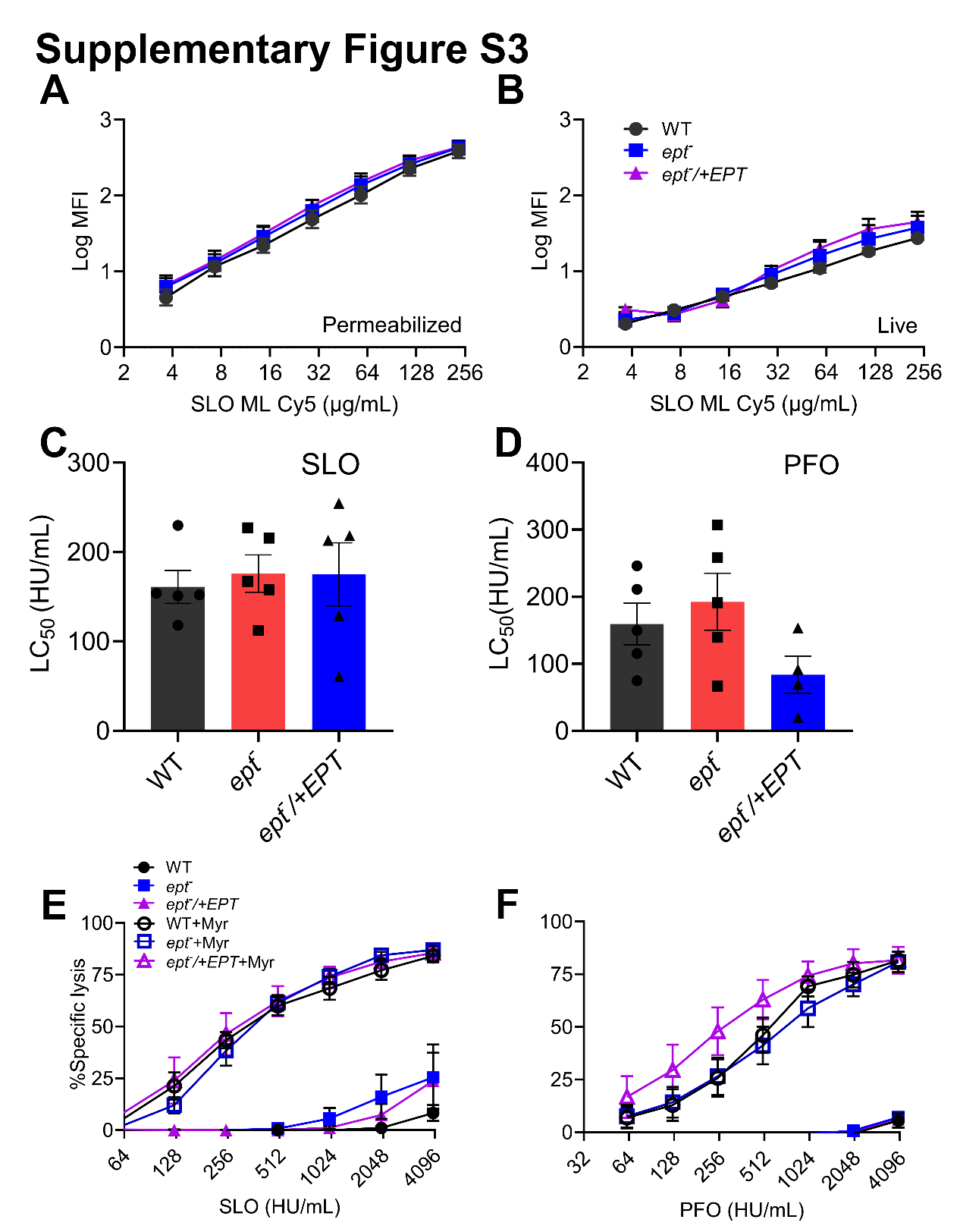

**Supplementary Figure** **S3.** **PME is dispensable for cytotoxicity and binding of CDCs to *L. major* promastigotes.** (A, B) Wild type (WT), *ept^—^*, or *ept^—^*/+EPT *L. major* promastigotes with or without 10 μM myriocin pretreatment were challenged with monomer-locked SLO conjugated to Cy5 at 4° C. Log median fluorescence intensity of Cy5 fluorescence gated on live and permeabilized cells are shown. (C-F) *L. major* promastigotes with or without 10 μM myriocin pretreatment were challenged with SLO (C, E) or PFO (D, F) at 37° C. PI uptake was measured by flow cytometry. (E, F) The LC_50_ values were calculated via logistic modeling. Graphs display mean ± SEM of (A, B) three or (C-F) five independent experiments, with (E, F) independent experiments plotted as individual points. Statistical significance was tested using 2 way ANOVA with multiple comparison and Sidak-Bonferroni correction.

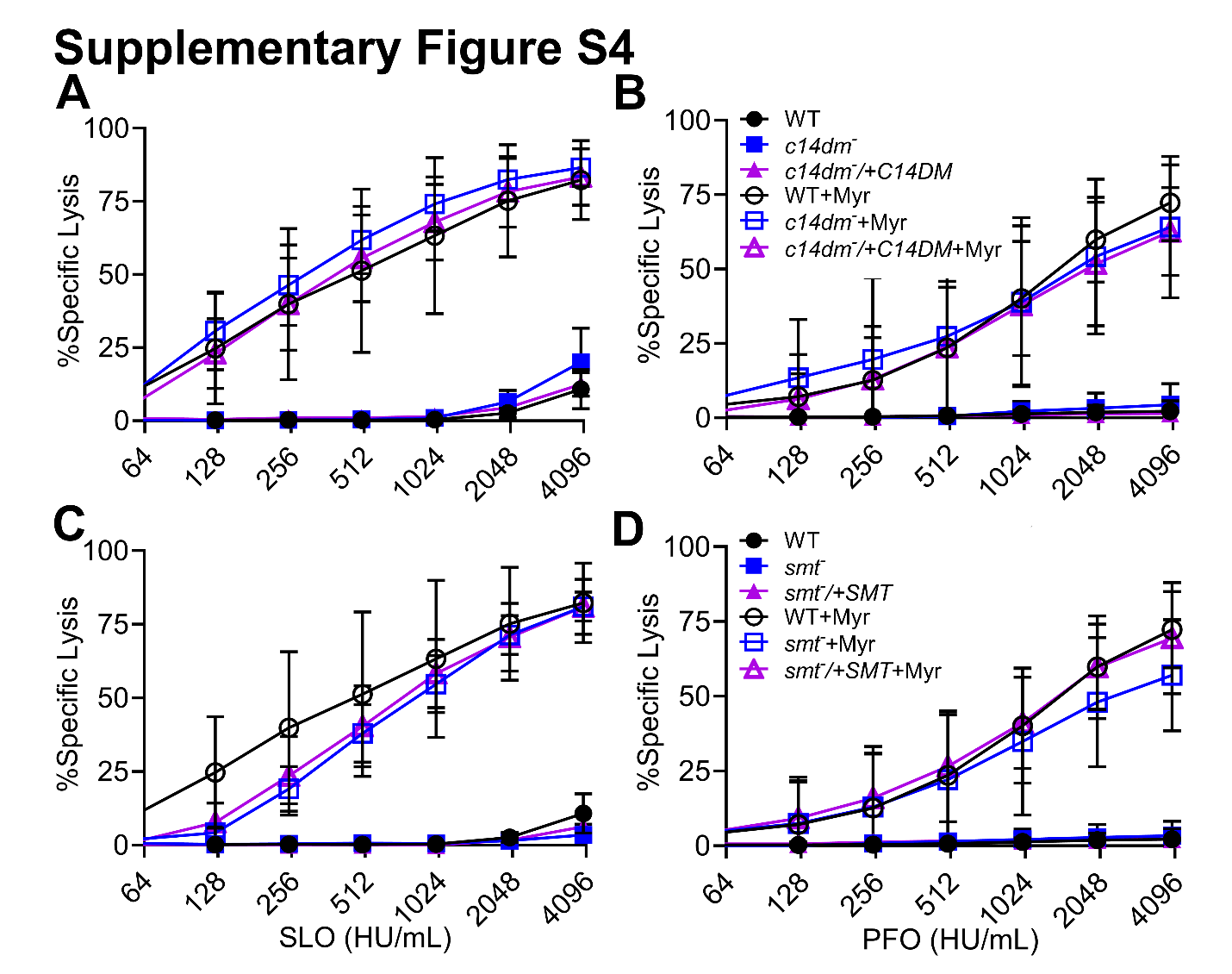

**Supplementary Figure** **S4. Ergosterol mutant *c14dm*^—^ and *smt*^—^ show sensitivity to CDCs only on myriocin treatment.** Wild type (WT), (A,B) *c14dm^—^*, *c14dm^—^*/+C14DM, (C, D) *smt^—^* , or *smt^—^*/+SMT promastigotes pretreated with vehicle or 10 μM myriocin were challenged with SLO or PFO at 37° C for 30 min at indicated concentrations. PI uptake was measured by flow cytometry. All genotypes were assayed together so the WT results are the same in A and C and B and D. Graphs are split for clarity. Graphs display mean ± SEM of three independent experiments.

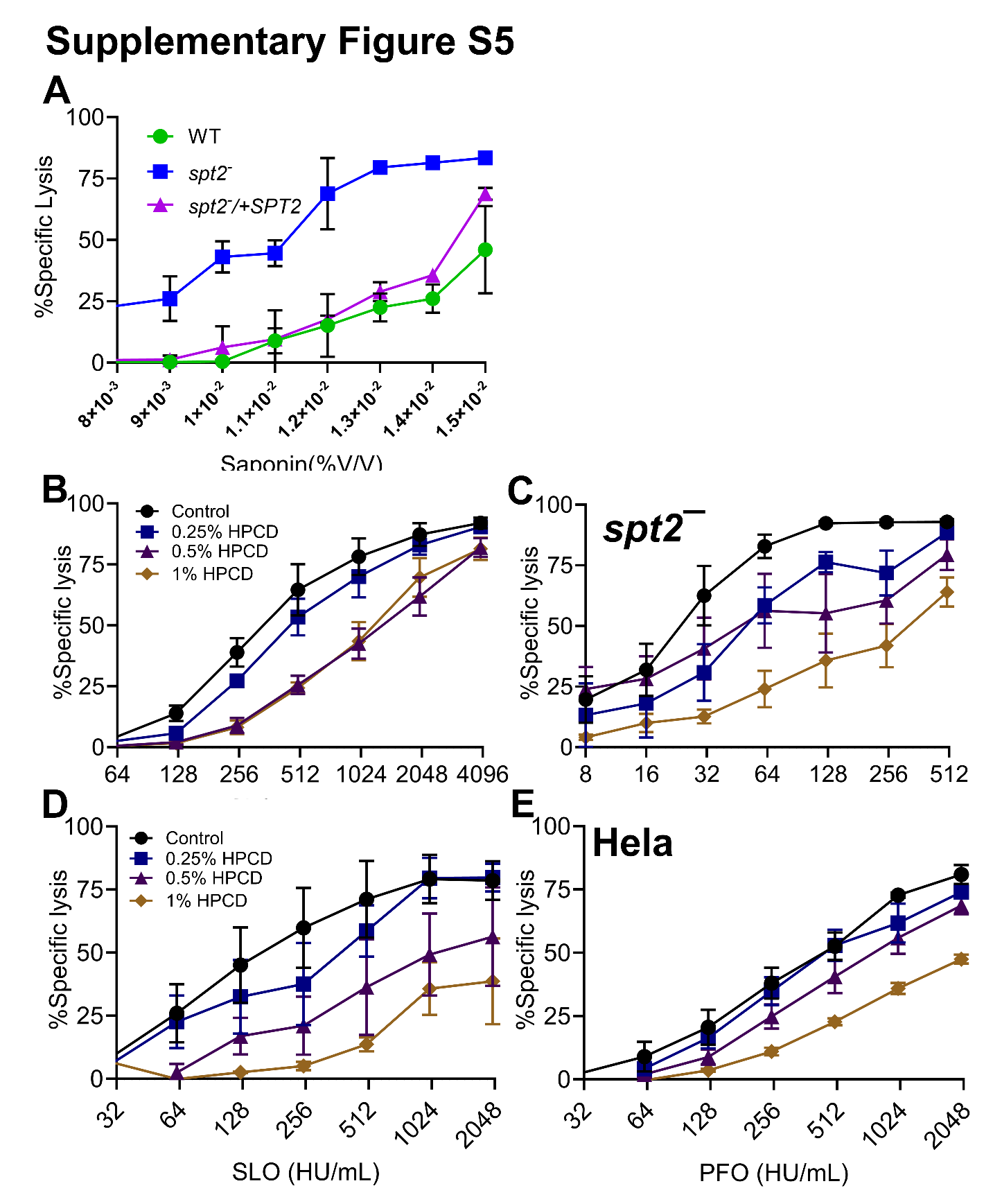

**Supplementary Figure** **S5. CDCs use sterols to lyse *L. major*.** (A) Wild type (WT), *spt2^—^*, and *spt2^—^*/+SPT2 promastigotes were challenged with indicated concentration of saponin (x 10^-2^ %V/V) at 37^ο^ C. PI uptake was analyzed by flow cytometry. (B, C) *L. major spt2^—^* or (D, E) Hela cells pretreated with indicated concentrations of 3-hydroxylpropylcyclodextrin (HPCD) were challenged with (B,D) SLO or (C,E) PFO at 37° C for 30 min. PI uptake was measured by flow cytometry. Graphs display mean ± SEM of three independent experiments.
